# Temperature dependence of platelet metabolism

**DOI:** 10.1101/802660

**Authors:** F. Jóhannsson, J.T. Yurkovich, S. Guðmundsson, Ó. E. Sigurjónsson, Ó. Rolfsson

## Abstract

Temperature plays a fundamental role in biology, influencing cellular function, affecting chemical reaction rates, molecular structures, and interactions. While the temperature dependence of many biochemical reactions is well defined *in vitro*, the effect of temperature on metabolic function at the network level is not well understood but remains an important challenge in optimizing the storage of cells and tissues at lower temperatures. Here, we have used time-course metabolomics data and systems biology approaches to characterize the effects of storage temperature on human platelets (PLTs) in platelet additive solution. We observed that changes to the metabolome with storage time do not simply scale with temperature but instead display complex temperature dependence, with only a small subset of metabolites following an Arrhenius-type. Investigation of PLT energy metabolism through integration with computational modeling revealed that oxidative metabolism is more sensitive to temperature changes than is glycolysis. The increased contribution of glycolysis to ATP turnover at lower temperature indicates a stronger glycolytic phenotype with decreasing storage temperature. More broadly, these results demonstrate that the temperature dependence of the PLT metabolic network is not uniform, suggesting that efforts to improve the health of stored PLTs could be targeted at specific pathways.

**Statement of Significance:** The temperature dependence of cellular metabolism is difficult to study due to regulatory events that are activated upon deviation from the optimal temperature range. Platelets are blood components used in transfusion medicine but also serve as a model cell to study human energy metabolism in the absence of genetic regulation. We investigated changes in platelet metabolism at temperatures spanning from 4 °C-37 °C using a quantitative metabolic systems biology approach as opposed to assessing individual reactions. We found that energy producing metabolic pathways have different temperature sensitivities. The results define the metabolic response to temperature on the metabolic pathway level and are of importance for understanding the cryopreservation of human platelets and more complex human cells used in cellular therapy.

## Introduction

Temperature affects many enzyme-catalyzed biochemical reactions and biological processes. In the late 19th century, van’t Hoff and Arrhenius (1) determined that the rate constant *k* for a large number of chemical reactions can be expressed as an explicit function of temperature

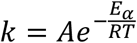

where *R* is the gas constant, *T* is absolute temperature, *E*_*α*_ is the activation energy of the reaction and *A* is known as the pre-exponential factor of the reaction. While this empirical relationship describes the reaction rate of a single chemical reaction, previous studies have shown that the rate of many other more complex processes—such as the chirping of tree crickets (2), muscle contraction (3), drug shelf life (4), and cellular and whole organism metabolism (5-7)—can be modeled by this equation. The rates of these phenomena all depend at least in part on chemical reaction rates. The temperature dependence of rates of biological processes is expressed in the unitless coefficient *Q*_*10*_, the ratio of reaction rates occurring at temperatures differing by 10°C (8). The van’t Hoff rule states that the *Q*_*10*_ for biological processes is generally between 2-3 (8-13), with some notable exceptions (6, 8). It is important to note that the *Q*_*10*_ is dependent on the measured temperature range, and its value decreases with increasing temperature (8, 9). However, it has been demonstrated that for certain biological processes, *Q*_*10*_ remains relatively stable over a range of temperatures (7, 8). Understanding the temperature dependence of biochemical reactions is fundamental to translating changes on the molecular level to higher levels of biological organization. Even with simple biochemical pathways, and in the absence of allosteric regulation, the pathway *Q*_*10*_ cannot be inferred from the temperature dependencies of its reactions (10). To bridge the gap between individual reactions and the whole metabolic network, metabolic modeling can be employed to predict fluxes of the entire network at different temperatures.

We previously integrated targeted time-course metabolomics data with a cell-scale metabolic model (14, 15) to characterize the temperature dependence of the human red blood cell (RBC) metabolic network (7). RBCs are an advantageous system to study due to the relatively small size of the metabolic network, lack of organelles, and lack of transcriptional and translational regulation. Human platelets (PLTs) lack a nucleus but contain mRNAs and the protein machinery necessary for *de novo* protein synthesis (16); however, the basal protein synthesis in PLTs is low and is mostly linked to signal-dependent activation (16, 17). Temperature-dependent changes in metabolic activity of resting PLTs are therefore unlikely to be heavily influenced by genomic regulation. While RBCs rely entirely on glycolysis for energy generation, PLTs contain mitochondria and therefore depend on oxidative phosphorylation to generate energy (18). Thus, the human PLT represents a good human cell model in which to study the network-level temperature dependence of oxidative cellular metabolism.

PLTs stored in a blood bank setting also represent a practical context for studying the temperature dependence of metabolism. Platelet concentrates (PCs) are used in modern medicine to either reduce the risk of bleeding or to control active bleeding in patients (19). Under current blood banking practices, PCs are stored in gas permeable plastic bags at 22°C under constant agitation. Storing platelets at such a high temperature limits the shelf life to 5-7 days (depending on the country), mainly due to the risk of bacterial infection and the deterioration of PLT quality over storage (the “platelet storage lesion” (PLS) (20)). Storing PCs at lower temperatures thus could provide increased storage time and reduced risk of bacterial infection. However, as first demonstrated by Murphy and Gardner in 1969, PLTs stored at 4°C show reduced post-transfusion survival compared with platelets stored at room temperature (21, 22). Nonetheless, there has been a growing interest in using cold stored PLTs to control bleeding in trauma patients due to superior hemostatic function and the immediate rather than long term needs of those patients (23, 24).

The metabolism of stored PLTs has been extensively studied (18, 25, 26) (27-30). Recently, we conducted a systems analysis of PLT metabolism during storage in PAS (30) using a modified version of a previously published PLT metabolic network reconstruction (31). During storage, PLTs can utilize a variety of nutrients (18) and rely on both glycolytic and oxidative metabolic pathways for energy production (25, 30). The temperature effects on platelet metabolism have mostly been investigated in the context of the glycolytic activity of PCs stored at 4°C by measuring the uptake and secretion rates of glucose and lactate, respectively (32-34). To the best of our knowledge, a formal quantitative study on the effects of temperature on the metabolic network has not been conducted. Here, we used targeted time-course metabolomics analysis of PCs stored to assess the PLT metabolic network at four different temperatures (4°C, 13°C, 22°C and 37°C). We integrated these data into a cell-scale computational model of PLT metabolism (15, 30) to predict the metabolic flux of the entire metabolic network, allowing for the estimation of *Q*_*10*_ values for individual metabolic reactions and pathways comparison of network-level responses to temperature changes.

## Methods

### Preparation and storage of PCs

Buffy-coat pools were obtained from 8 healthy donors Blood Bank, Landspitali-University Hospital, Iceland. Four leukoreduced buffy-coat pools were combined and then split into four identical PCs containing approximately 65% platelet additive solution (SSP+, MacoPharma) and 35% plasma. The PCs were stored in plastic containers (PL 2410, Fenwal) under constant agitation at four different temperatures: 4°C, 13°C, 22°C, and 37°C. The National Bioethics Committee of Iceland approved the study.

### Sample collection

Samples were collected at seven different time points from each PC. From the PC stored at 4°C samples were collected at 24h, 48h, 108h, 216h, 324h, 432h, and 552h; from the PC stored at 13°C samples were collected at 24h, 48h, 60h, 120h, 180h, 240h, and 312h; from the PC stored at 22°C samples were collected at 24h, 48h, 72h, 96h, 120h, 144, and 168h; and from the PC stored at 37°C 3h, 6h, 24h, 27h, 30h, 48h, and 51h. Samples were collected as previously reported (28).

### Assays

Immediately after sample collection, a hematology analyzer (CELL-DYN Ruby, Abbott Diagnostics, Abbott) was used to measure the platelet count, and a blood gas analyzer (ABL90 FLEX, Radiometer MedicalApS, Brønshøj, Denmark) was used to measure the concentrations of glucose and lactate. The extracellular acetate concentration was measured with an enzymatic coupled assay (ACETAK), according to the manufacturer’s instruction (Megazyme International Ireland Ltd, Wicklow, Ireland) using a microplate reader (Spectramax M3, Molecular Devices, Sunnyvale, CA).

### Metabolomics

The metabolomics analysis follows the same procedure as explained in (28). Briefly: Samples were split into extra- and intracellular fractions by centrifugation. Polar metabolites were extracted using cold methanol:water extraction (7:3) followed by evaporation and resuspension in acetonitrile:water (1:1). The metabolites were separated and detected using ultra-performance liquid chromatography (UPLC; Acquity, Manchester, UK) coupled to a quadrupole-Time of flight mass spectrometry (Q-TOF MS; Synapt G2, Waters). Chromatographic separation was achieved by hydrophilic interaction liquid chromatography (HILIC) using two conditions: 1) Acidic mobile phase (ACN:water w. 0.01% formic acid) measured in both positive and negative ionization modes. 2) Basic mobile phase (ACN:water w. 10 mM ammonium bicarbonate) measured in negative ionization mode. A calibration mixture of 72 metabolites was measured at eight concentrations within each batch. All samples were measured in triplicates in a randomized order. The metabolites were identified by matching chromatographic retention times and mass-to-charge ratios to our in-house database. For the relative quantification of targeted metabolites Targetlynx (v4.1, Waters) was used to integrate the chromatographic peaks which were normalized to appropriate internal standards. Absolute quantifications of selected metabolites were carried out by using standard curves generated by the calibration mixture. Principal components analysis (PCA) was performed using R v3.5.1 (R Development Core Team, Vienna, Austria). Prior to the PCA the data were normalized using z-score normalization. Hierarchical clustering and heatmap generation were performed using MetaboAnalyst v.3 (35).

### Temperature coefficient Q_10_ calculation

The *Q*_*10*_ coefficient describes the ratio of rates between temperatures differing by 10°C. The *Q*_*10*_ is given by

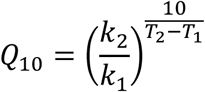

where *T*_*1*_ and *T*_*2*_ are temperatures in Kelvin, *T*_*2*_ > *T*_*1*_, and *k*_*1*_ is the rate at *T*_*1*_ and *k*_*2*_ the rate at *T*_*2*_. In cases where *Q*_*10*_ is relatively temperature invariant, a regression line can be fitted to the logarithm of the reaction rates against temperature according to

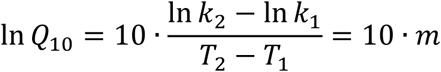

where the slope of the regression line *m* can then be used to estimate *Q*_*10*_ through

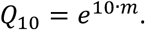

### Constraint-based metabolic modeling

To estimate the flux states at each temperature, we used a modified version of the platelet metabolic reconstruction iAT-PLT-636 (31) following the same procedures as in (30). This modified version has previously been used for metabolic network analysis of buffy coat and apheresis derived PCs stored in PAS (30). Here, further modifications have been made to accommodate choline metabolism by adding the choline kinase reaction (CHOLK):

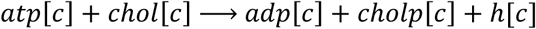

This reaction can be catalyzed by the choline/ethanolamine kinase *CHKB*, which has been detected in platelets by proteomics analysis (30). For each temperature, and each extracellular metabolite used to constrain the model, a regression line was fitted to the time-concentration profile. The slopes of the regression lines, normalized to the average platelet count at each condition, were used to estimate the uptake/secretion rate of the selected metabolites, similar to previous studies (15). In the case of infeasible models, an optimization algorithm was used to slightly adjust the constraints in order to obtain a feasible solution, as described in (30). For each constrained model, random sampling of the solution space (36) was carried out 5000 times. The model analysis was carried out in the MATLAB environment (Mathworks, Natick, MA, USA) using the COBRA Toolbox (37).

## Results

### Metabolite time-concentration profiles scale non-uniformly with temperature

We analyzed both extra- and intracellular metabolites in stored PCs. A total of 62 metabolites were measured in the extracellular fractions and 72 metabolites in the intracellular fractions. Underlying trends in the metabolomics data were analyzed by principal component analysis (PCA) (38) to identify how the metabolome scales with temperature. We observed a clear separation in both extra- and intra-cellular metabolic profiles of the PCs stored at different temperatures (Figure 1). Notably, we observed that the largest variance was due to storage time in the extracellular fraction (Figure 1a) but storage temperature in the intracellular fraction, with increasing separation with increased storage time (Figure 1b). The extra- and intra-cellular metabolic profiles were analyzed by hierarchical clustering with a heat map to visualize the time-dependent effects temperature has on the metabolome. In the extracellular fractions, distinct groups of metabolites were observed (Figure 2). A group of metabolites increased in concentration with storage time at all temperatures with rates proportional to temperature (including phenylalanine and xanthine). Another group (including cysteinylglycine disulfide and cysteineglutathione disulfide) decreased in concentration with rates inversely proportional to temperature. In other words, the rate of formation and breakdown of a subset of the extracellular metabolome increased with increased temperature. However, this trend was not observed for all measure metabolites; for example, the concentrations of aspartate and taurine increased with storage time at rates inversely proportional to temperature, while some metabolites were only secreted at 37°C (e.g., sphingosine 1-phosphate (S1P) and acetylcarnitine).

**Figure 1.**
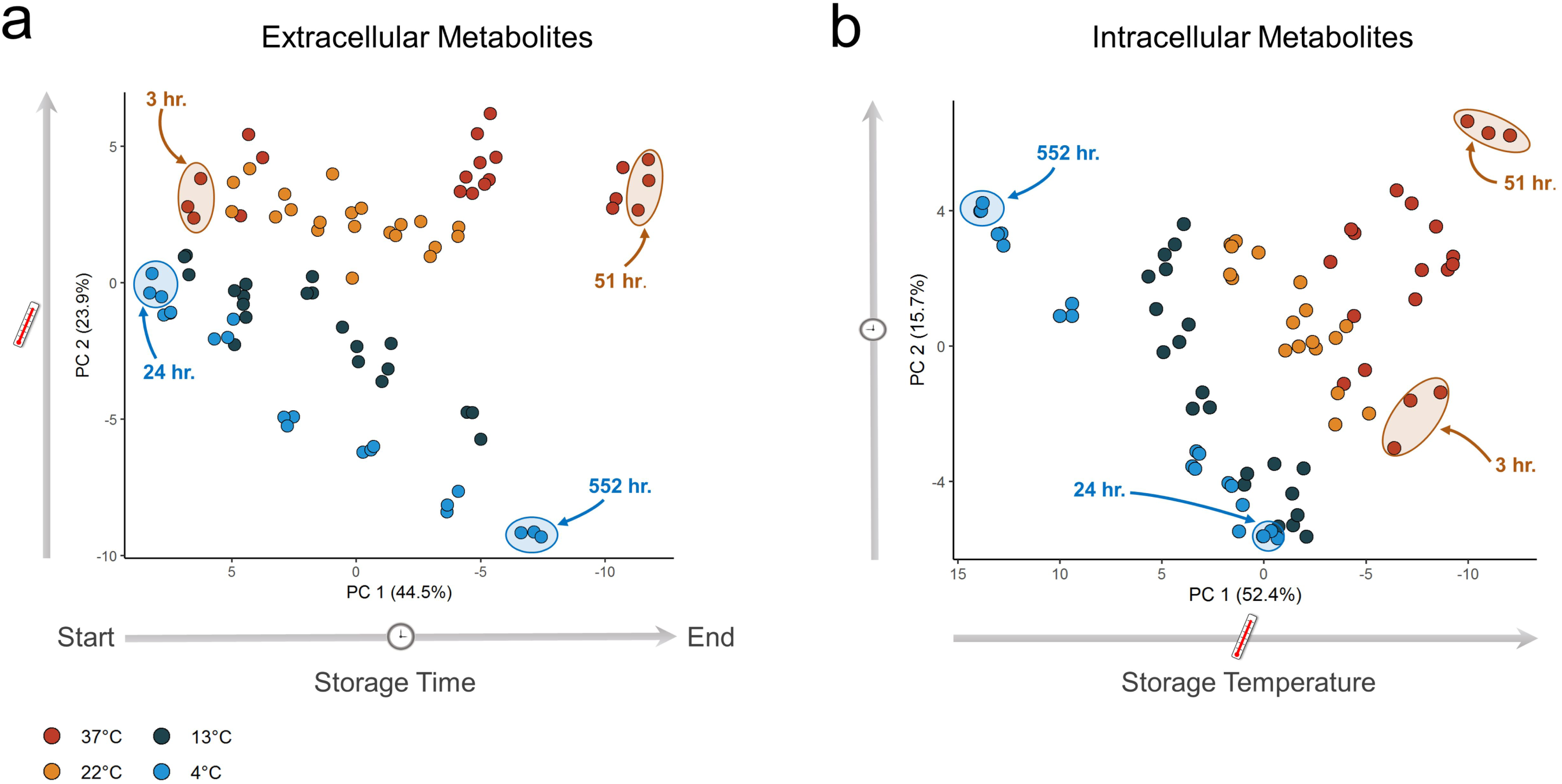
Principal component analysis (PCA) of extra- and intracellular metabolite data. Samples were collected at seven time points at four different temperatures, all samples were measured in triplicates. **a** PCA scores plot for the extracellular fractions. **b** PCA scores plot for the intracellular fractions.

**Figure 2.**
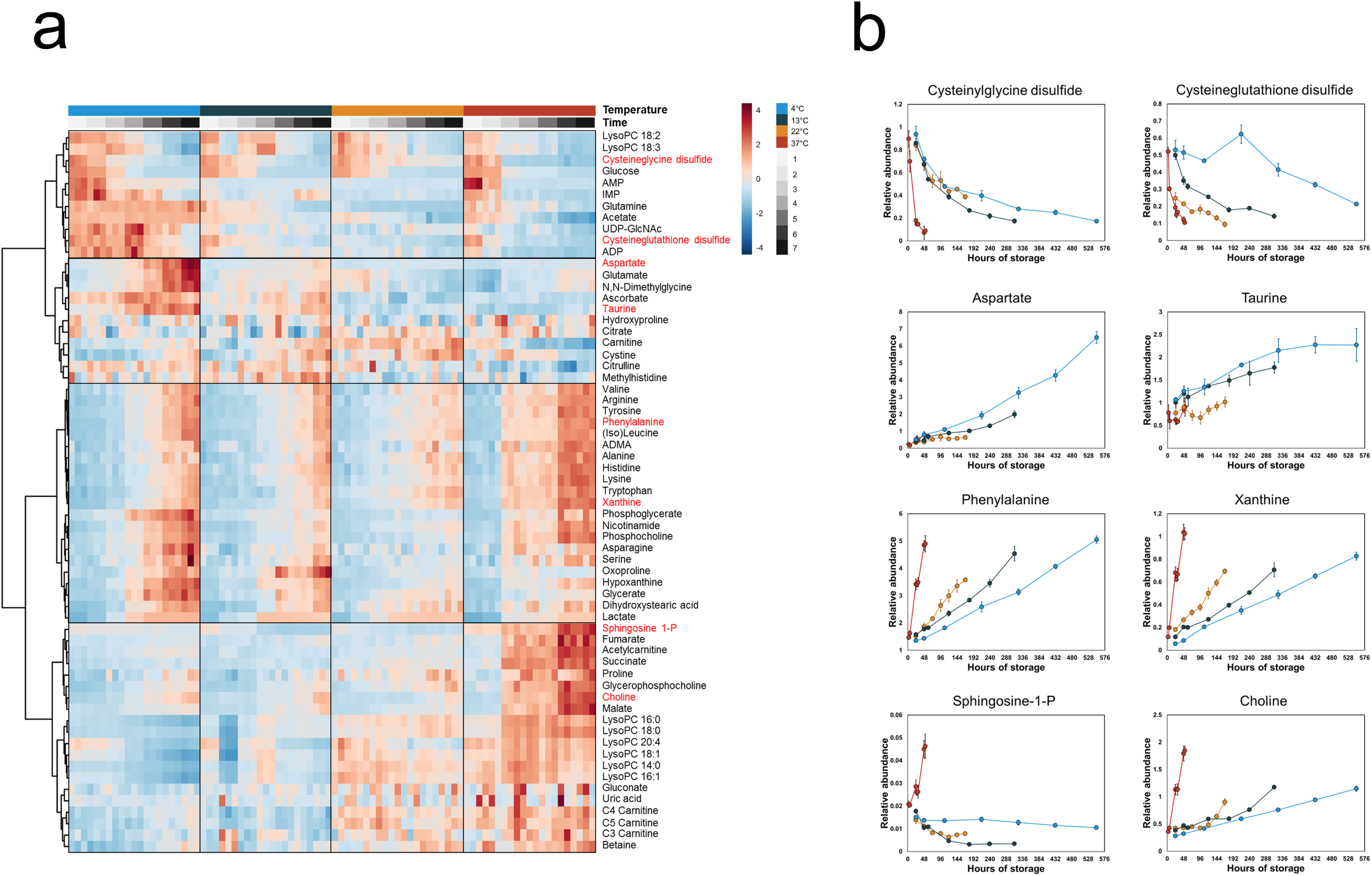
Time-concentration profiles of the extracellular metabolites. **a** A heatmap of all measured extracellular metabolites. Hierarchical clustering grouped the metabolites into four distinct clusters. **b** Line plots of selected metabolites representative of each cluster. The points represent the mean of the normalized metabolite abundance and the error bars standard deviation.

In the intracellular fractions (Figure 3), we observed higher concentrations of most measured metabolites at higher temperatures. A large fraction of the metabolome increased with the highest rate in the 37°C PLTs, including many intermediates of phosphatidylcholine degradation (e.g., lysophosphatidylcholines (LysoPCs) and choline). Conversely, a group of metabolites — including glutathione and S1P — were maintained at higher concentrations at 4°C compared to the other temperatures. In the PLTs stored at 4°C, there was a noticeable drop in concentration for most of the measured metabolites at 324h with the notable exception of adenine and adenosine. These metabolites increased sharply in concentration at the same time point, coinciding with the depletion of extracellular glucose. These results demonstrate that the temperature-dependence of the metabolome of stored PLTs is complex, that is, the reaction rates, do not uniformly increase with increased temperature.

**Figure 3.**
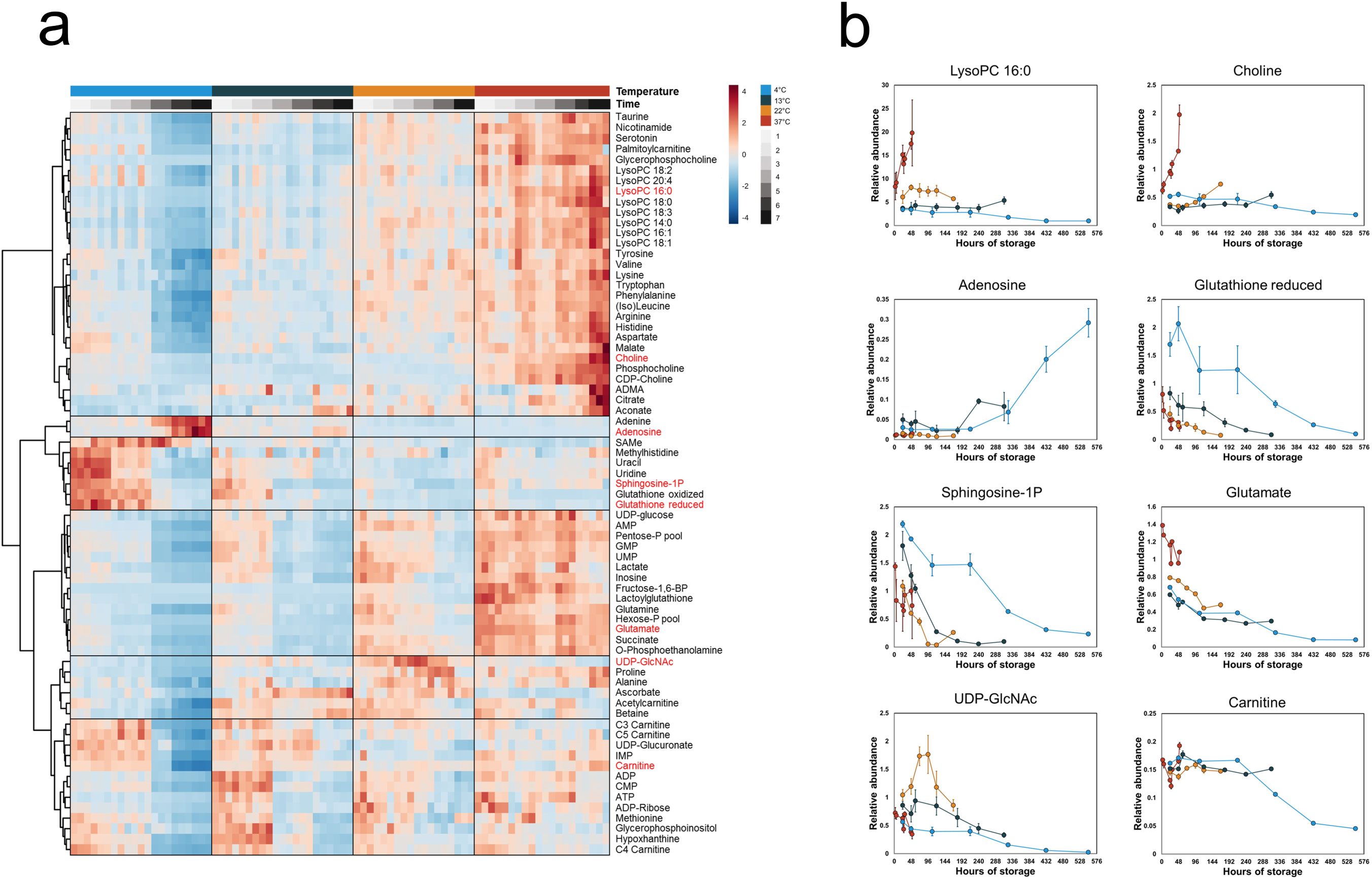
Time-concentration profiles of the intracellular metabolites. **a** A heatmap of all measured intracellular metabolites. Hierarchical clustering grouped the metabolites into six distinct clusters. **b** Line plots of selected metabolites representative of each cluster. The points represent the mean of the normalized metabolite abundance and the error bars standard deviation.

### A subset of extracellular metabolites follows an Arrhenius-type relationship with temperature

The data summarized in Figure 1 demonstrate that platelet metabolism follows a temperature-specific trajectory, in contrast to what would be expected if the rate of change of the metabolome would simply scale uniformly with temperature. However, as observed in Figure 2, a subset of the metabolites did scale with temperature. The extracellular metabolites represent end nodes of the pathways that produce or consume them, acting as proxies for the temperature dependences of those pathways.

To estimate *Q*_*10*_ for the rate of change of concentration of the extracellular metabolites, only metabolites where the concentrations varied linearly (*R*^*2*^ > 0.7) with time during the entire storage time were selected. With this criterion, the concentration of 17 metabolites were found to vary linearly with storage time, and their log-transformed rates varied linearly with temperature, with *R*^*2*^ ranging from 0.80 to 0.99 (Table 1). This linearity allowed for a single *Q*_*10*_ estimation for the entire temperature range. The estimated *Q*_*10*_ values for the extracellular metabolites ranged from 1.54 to 3.04 with a median of 2.15 (Table 1), a finding in good agreement with the expected range of 2–3 for *in vitro* biological reactions (7).

**Table 1.**
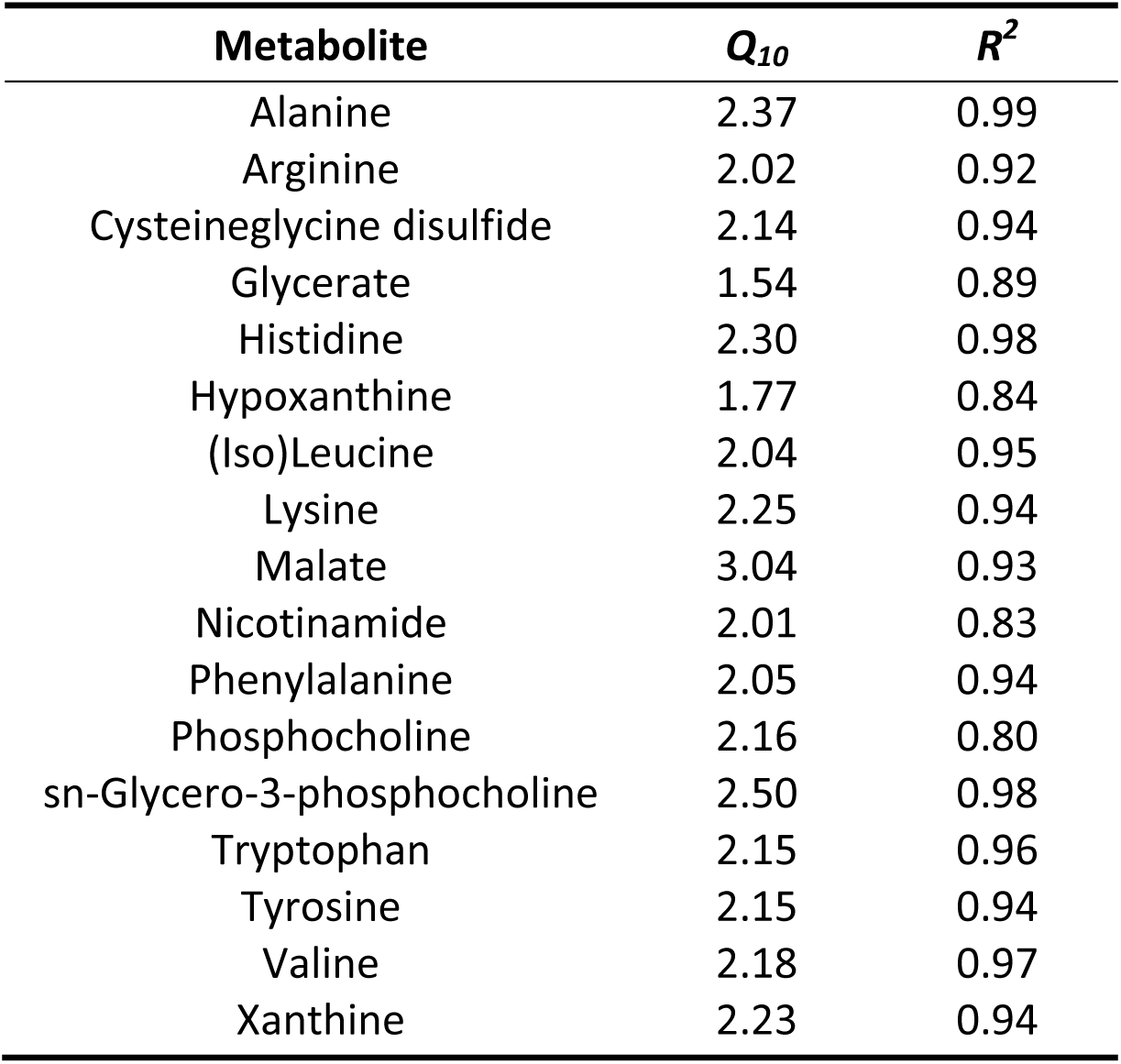
Q_10_ coefficient for extracellular metabolites.

### Acetate and glutamine metabolism is more sensitive to temperature than glucose metabolism

The most important fuels for ATP turnover in PCs stored in PAS at standard storing conditions (22°C) are in order: acetate, glucose, and glutamine (30). The time-concentration profiles of these three metabolites and lactate increased at different rates with temperature (Figure 4). Acetate and glutamine uptake halted at 4°C, while changes to glucose uptake and lactate secretion rates were observed to be less sensitive to temperature. The changes in uptake and secretion rates were quantified by estimating the respective *Q*_*10*_ coefficients. Since no detectable uptake of acetate was recorded at 4°C, it was excluded from the *Q*_*10*_ calculation. Acetate uptake between temperatures 13–37°C had a calculated *Q*_*10*_ = 2.98, which was relatively temperature invariant (*R*^*2*^ > 0.99). Glutamine uptake (Figure 4) occurred at similar rates at 37°C and 22°C but decreased at 13°C. Glutamine uptake did not display an Arrhenius-type relationship with temperature, and therefore, the *Q*_*10*_ was not estimated. In all PCs, glucose was depleted during the recorded storage time; only time points with non-zero concentrations of glucose and lactate were used for the *Q*_*10*_ calculations. We calculated lower *Q*_*10*_ values for glucose (1.73, *R*^*2*^ > 0.99) and lactate (1.70, *R*^*2*^ = 0.98). The differences in the time-concentration profiles between 13°C and 22°C were not substantial (Figure 4). Nevertheless, changes to temperature influenced the uptake and secretion rates of glucose and lactate to a lesser degree than acetate and glutamine. In addition, their uptake did not halt at 4°C as observed for acetate and glutamine. In PLTs, glucose is primarily metabolized via glycolysis while acetate and glutamine fuel the TCA cycle. These results reflect an altered scaling of metabolic pathways in stored platelets with temperature.

**Figure 4.**
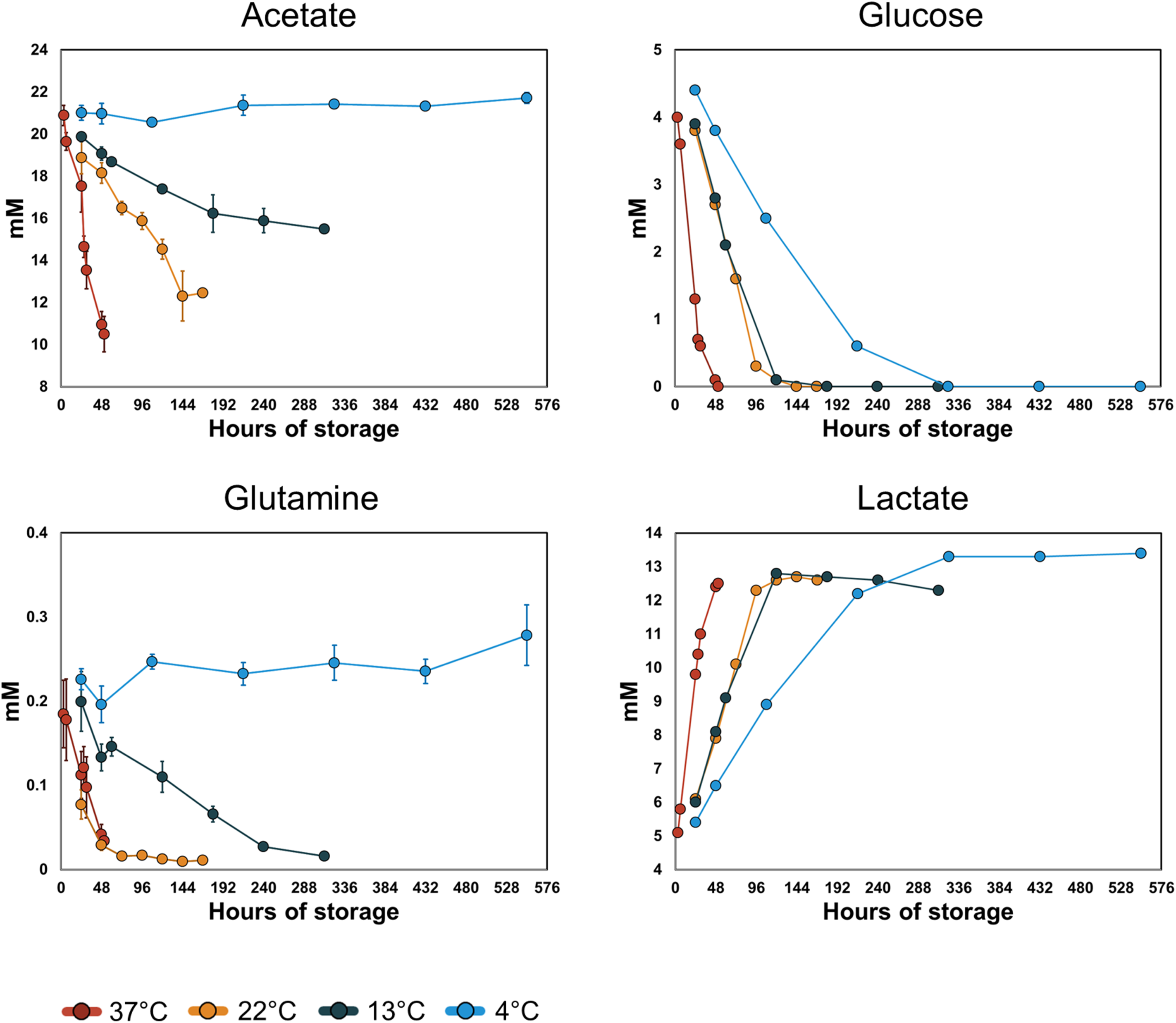
Time-concentration profiles of the main energy producing metabolites and lactate. The points represent the mean of the measured value and the error bars the standard deviation. The rates of uptake of acetate and glutamine increase with increasing temperature with no detected uptake at 4°C. The rates of glucose uptake and lactate secretion increase with increasing temperature. In all PCs glucose is depleted during the recorded storage time. Due to this only time points were glucose and lactate concentrations with non-zero concentrations of glucose and lactate were used for the *Q*_*10*_ calculations.

### Metabolic modeling reveals pathway specific temperature dependence

In order to investigate the temperature dependence on the metabolic pathway level, we integrated the measured metabolomics data with a metabolic network reconstruction of the human PLT (30, 31). Previous metabolomics studies on PLTs stored in PAS revealed discreet metabolic phenotypes over storage time, with a transition from the first to second phenotypes occurring at day 4 of storage (28, 29). For the metabolic modeling, only the first phenotype (days 1-3) was chosen because *(i)* the first phenotype represents relatively fresh PLTs that still have not gone through extensive PSL, and (*ii*) at this stage the extracellular glucose had not been depleted, and its uptake rate was relatively constant.

To estimate the equivalent time period for the PCs stored at temperatures other than 22°C, relative aging was assumed to follow an Arrhenius model. A *Q*_*10*_ of 2.0 was chosen roughly corresponding to time points 3-24h at temperature 37°C, 24–72h at 22°C, 24–120h at 13°C and 24–216h at 4°C. At these time points, glucose has not been depleted and the rates of change in concentration vary linearly with time for glucose, lactate, and acetate (except at 4°C). To estimate *Q*_*10*_ coefficients for individual reactions the following criteria were used: only reactions active (i.e., rate > 10^−6^ mmol/day/PLT·10^12^) at all temperatures were used and only reactions where the *Q*_*10*_ coefficients were relatively temperature invariant (*R*^*2*^ > 0.7). A total of 579 reactions were predicted to be active at all temperatures, of which 94 reactions (excluding transporters) had relatively temperature-independent *Q*_*10*_ coefficients with a median of 2.23 (1.72–2.80). To investigate the relative temperature sensitivity of distinct metabolic pathways, the 94 reactions were sorted into 13 subsystems (Table 2) as defined by the global human metabolic network reconstruction (39). A notable difference in *Q*_*10*_ values was observed in the energy-producing subsystems: glycolysis, citric acid cycle and oxidative phosphorylation.

**Table 2.**
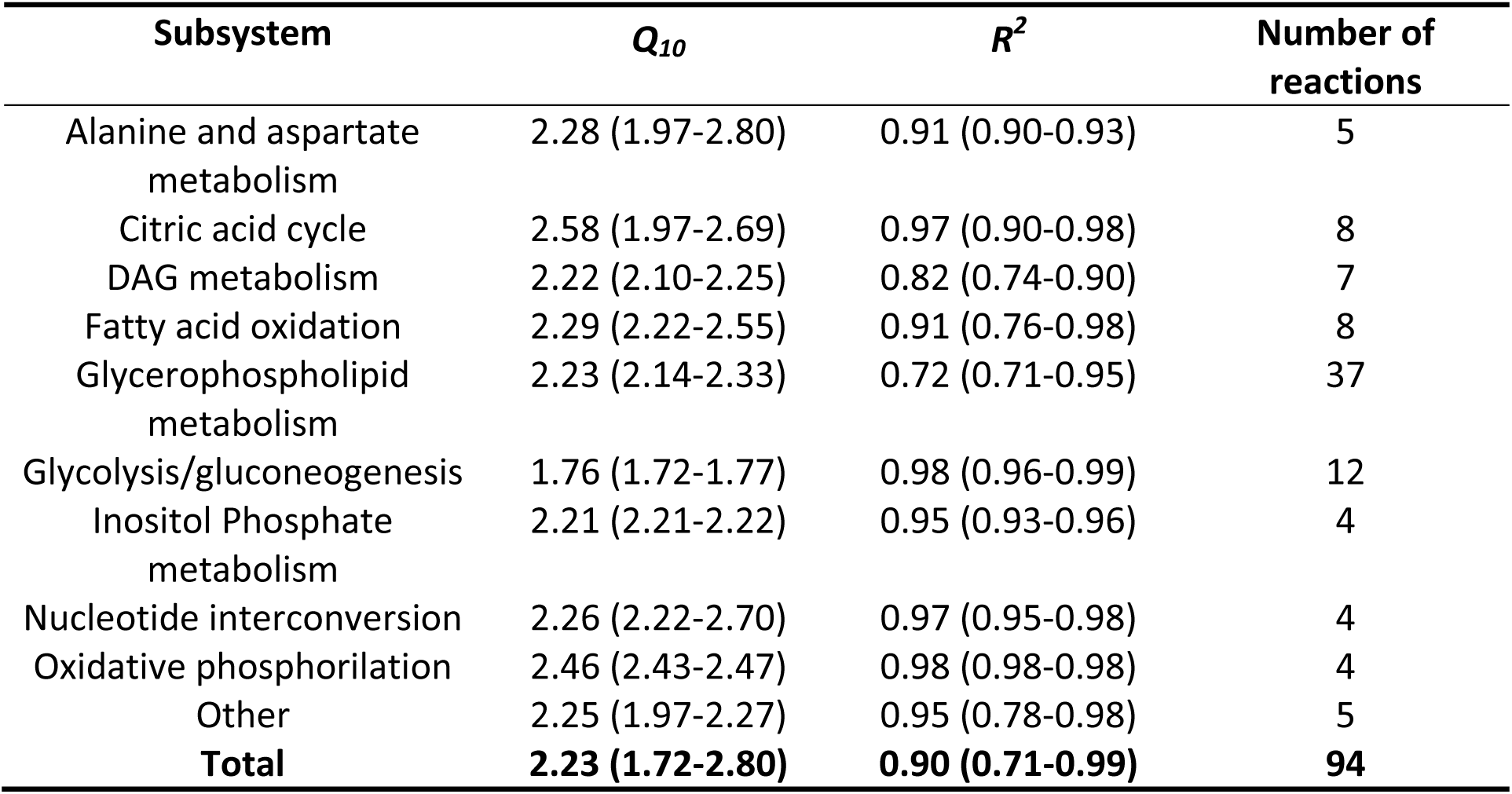
The distribution of predicted *Q*_*10*_ in different subsystems of the metabolic network. The table dispays median values with minimal and maximal values in parenthesis.

Constraint-based modeling offers a framework for analyzing metabolism at the cellular level (14, 40), allowing for quantitative predictions of cellular functions such as net ATP production and the relative contribution of the glycolytic and oxidative pathways to net ATP production (30). Constraint-based models have previously been used to estimate the temperature dependence of a metabolic network (7), suggesting that the temperature dependence of these cellular processes could also be computed through the integration of metabolomics data (15). The predicted *Q*_*10*_ for net ATP production was 2.24 (R^2^ 0.99). At all temperatures, oxidative metabolism was predicted to account for a higher proportion of ATP production. This proportion was, however, temperature dependent and increased with increasing temperature from 69.5% at 4°C to 86.2% at 37°C (Figure 5).

**Figure 5.**
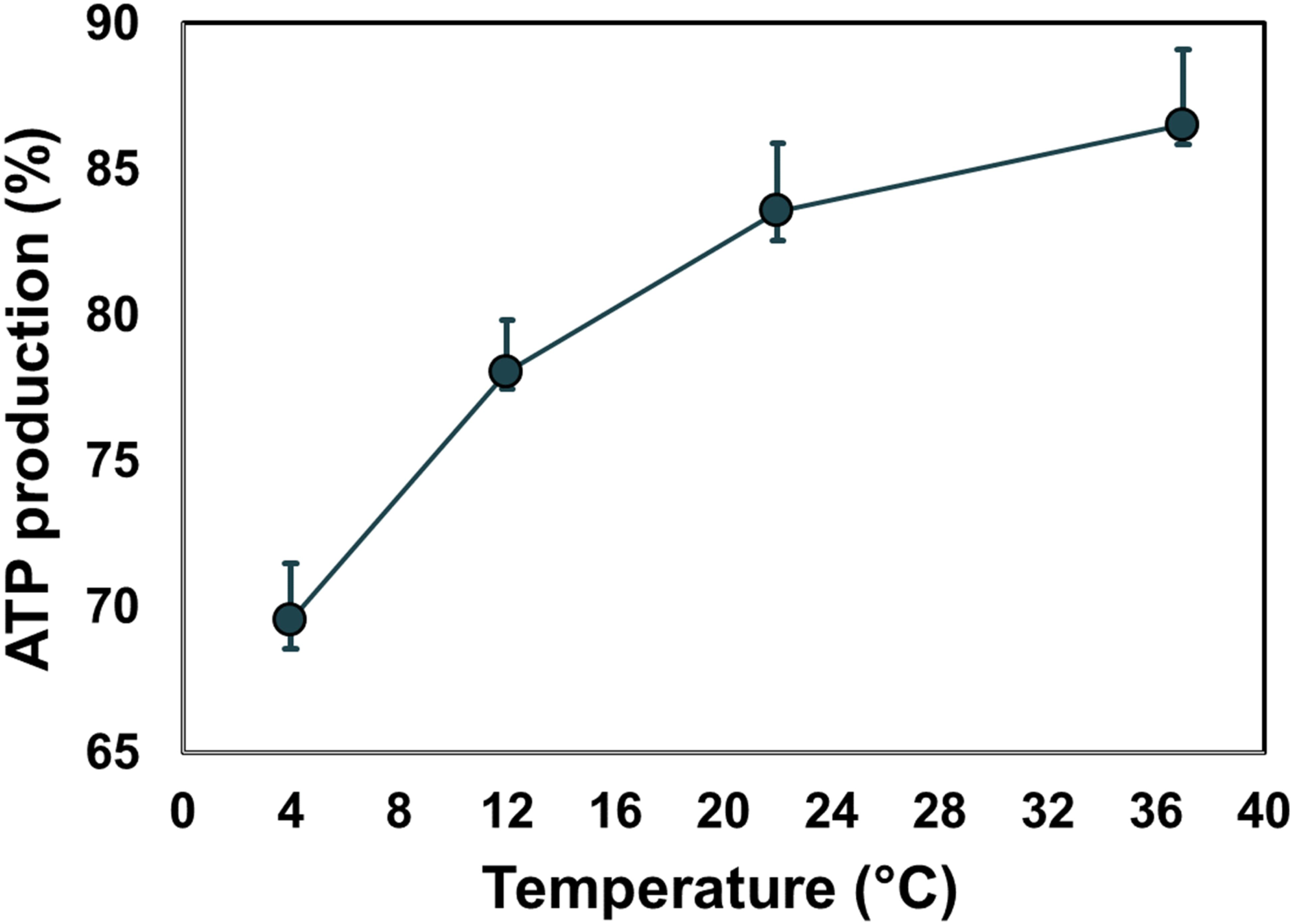
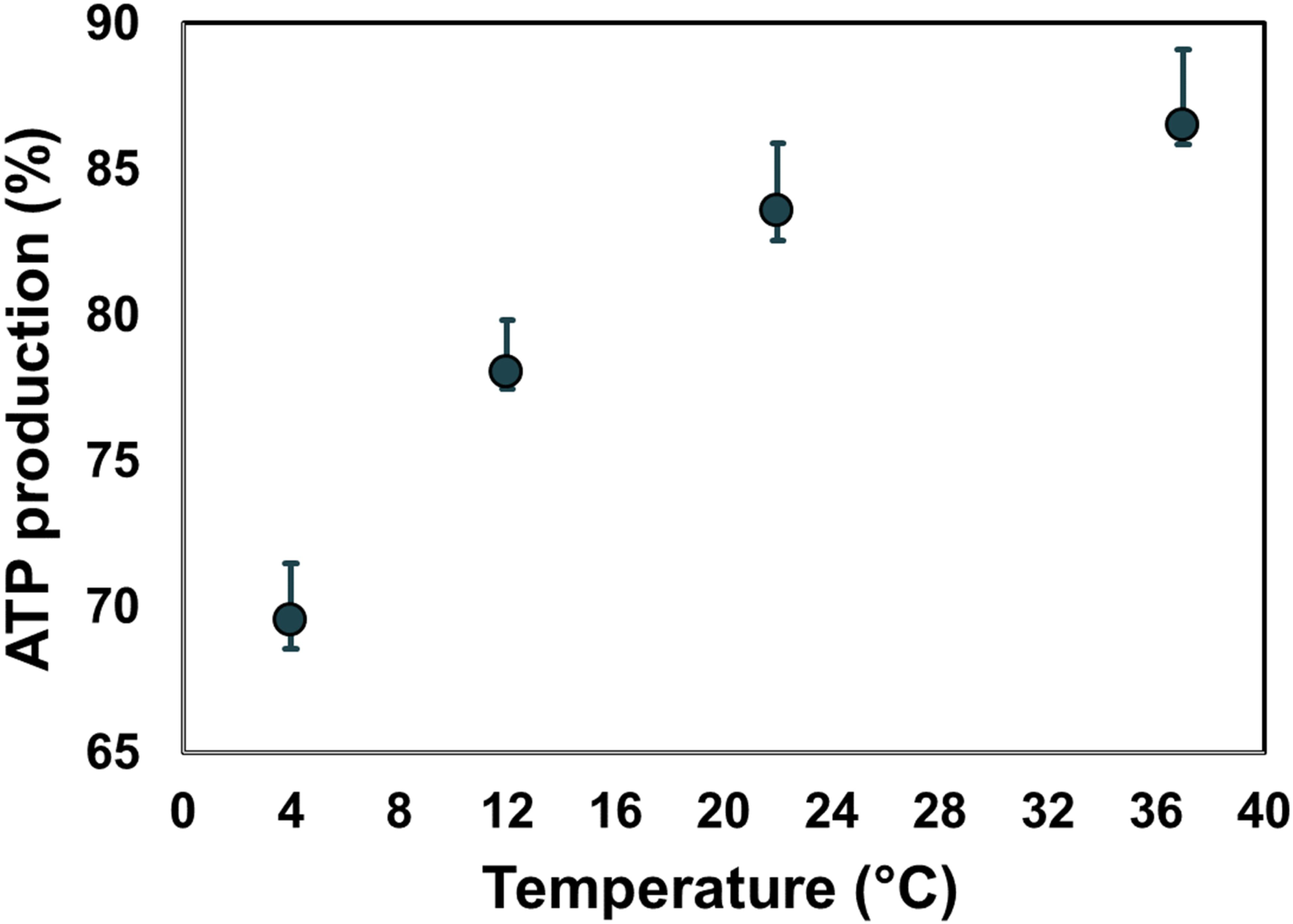
The relative contribution of oxidative metabolism to ATP production at different temperatures. The points represent the median of the random sampling results and the error bars the minimum and maximum.

## Discussion

Temperature influences the rates of biological processes. Here, we have investigated the temperature dependence of the PLT metabolic network during storage in PAS. We observed a non-uniform response over a 33°C temperature range for 134 measured metabolites. This result is unsurprising since the metabolome represents the output of a complex system that depends on the kinetic properties of a large number of enzymes, as well as many other factors such as membrane fluidity, ionic strengths and protein-protein interactions. Nonetheless, for a subset of the measured extracellular metabolites, the rate of change in concentration remained relatively constant throughout the storage time within each measured temperature. These rates could be modeled with an Arrhenius-type relationship to yield a single *Q*_*10*_ value over the entire temperature range. A global analysis of the metabolic network—using both metabolomics data and computational models—indicated that pathway usage shifts as a function of temperature and is different dependent upon the metabolic pathway.

Platelets are highly metabolically active cells (18) that utilize both glycolytic and oxidative metabolic pathways for energy production. Both processes are sensitive to temperature changes. The glycolytic activity can be assessed by measuring the uptake rate of glucose and secretion rate of lactate. The decrease in glycolytic activity at low temperatures reported here is in good agreement with previous studies comparing these parameters in PCs stored at 22°C and 4°C (32-34). Between temperatures 4-37°C these rates, and as a proxy the glycolytic rate, could be modeled by the Arrhenius equation with *Q*_*10*_ ≈ 1.7, meaning that for every 10°C increase in temperature glycolytic activity increases 1.7-fold. Oxidative metabolism provides the bulk of ATP turnover in platelets (18, 25, 30). In PCs stored in PAS at room temperature, acetate is the primary oxidative fuel and is mostly converted to CO_2_ via oxidative phosphorylation (27, 30). The *Q*_*10*_ coefficient for acetate uptake could not be estimated for the entire temperature range since no acetate uptake was detected at 4°C. Acetate uptake was measured over the temperature range 13-37°C, and a single value for the temperature coefficient could be obtained at *Q*_*10*_ = 2.98. These results indicate that oxidative metabolism of stored PLTs is more sensitive to temperature changes than the glycolytic metabolism over the 4-37°C range. Glutamine can be utilized as an oxidative fuel by platelets via glutaminolysis, which provides substrates for the TCA cycle (30, 41). The decrease in the uptake rate of glutamine at 13°C and the absence of net uptake at 4°C further indicate a slowdown of oxidative metabolism with decreasing temperature.

The PLT cell-scale model allows for predictions of flux distribution over the entire metabolic network given a set of constraints in the form of uptake/secretion rates. Here we have utilized the PLT model to predict flux state at four different temperatures by adding constraints of 20 extracellular metabolites measured at all temperatures. The results show that the *Q*_*10*_ values for stored PLTs fall within a relatively narrow range (1.72-2.80) compared to the previously computed flux states of stored RBCs (7). Unsurprisingly, the overall metabolic rate decreased with decreasing temperature. The decrease in flux is however, not uniform, so the relative fluxes through reactions and pathways changes with temperature.

Flux through metabolic pathways is controlled by many factors, including the kinetic properties of the constituent enzymes, concentration of reactants and products, and activity of other pathways and cellular processes. In *E.coli*, for example, ATP-demanding processes have been shown to have the largest influence on the glycolytic flux (42). In human PLTs, a decrease in oxidative metabolism either by inhibitors (41, 43) or hypoxia (44) can be, at least partially, compensated for by increased glycolytic flux. Furthermore, platelet activation—an energy demanding process—leads to increases in both glycolysis and oxidative metabolism (41, 43), meaning that under basal conditions these pathways are maintained below their maximum capacity. It is therefore reasonable to assume that ATP-demand is an important factor in the flux control of in PLTs, and that the thermal effects seen here on the PLT metabolic network are governed both by the temperature dependencies of the reactions within the pathways as well as the temperature dependencies of ATP-demanding processes. Thus, we hypothesize that the higher thermal sensitivity of oxidative metabolism leads to an upregulation in glycolysis, and consequently that the glycolytic rate at lower temperature is maintained closer to its maximum capacity, which would explain the relatively low *Q*_*10*_ of glycolysis compared to the RBCs (7) and other pathways in the PLTs.

The observed shift from oxidative metabolism may be beneficial to PLT storage. PLTs stored at 4°C have been demonstrated to have lower mitochondrial reactive oxygen species (ROS) levels, leading to a better preservation of mitochondrial function (45). This observation could explain the higher levels of glutathione at lower temperatures reported here (Figure 3B).

One possible mechanism for this phenomenon is a decrease in flux through the complexes of the electron transport chain; however, the rate of mitochondrial ROS generation depends not only on the oxygen consumption rate but also on other factors such as the protonmotive force, the rate constant of O^2•^ production and the mitochondrial NADH/NAD^+^ ratio (46, 47). Thus, the data presented here is not sufficient to provide a mechanistic link between temperature and ROS generation.

In this study, we have demonstrated that lowered temperature slows down oxidative phosphorylation more than glycolysis in stored PLTs. The fact that acetate and glutamine uptake are brought to a halt at 4°C puts it into question whether oxidative metabolism is active at all at low temperatures, and the need for oxidative fuel in the additive solution in cold stored platelets. Conversely, the relative contribution of glycolysis to energy production increases at lower temperatures necessitating ample extracellular glucose concentration for long term storage. A limitation of the study is that no direct measurements of oxygen uptake are presented, only predicted values based on the constraint-based modeling. Our results demonstrate that the response of the PLT metabolic network to temperature change is complex and causes shifts in relative pathway activity, most notably a shift from oxidative to glycolytic metabolism at lower temperatures.

## Author contributions

F.J. and O.E.S. performed the experiments. F.J. analyzed the data and wrote the manuscript. J.Y and S.G. contributed to data analysis and to writing of the manuscript. O.R. and O.E.S. devised the study and contributed to writing of the manuscript.

## Acknowledgments

The authors would like to express their thanks to the staff at the Icelandic Blood Bank.

